# DandD: efficient measurement of sequence growth and similarity

**DOI:** 10.1101/2023.02.02.526837

**Authors:** Jessica K. Bonnie, Omar Ahmed, Ben Langmead

## Abstract

Genome assembly databases are growing rapidly. The sequence content in each new assembly can be largely redundant with previous ones, but this is neither conceptually nor algorithmically easy to measure. We propose new methods and a new tool called DandD that addresses the question of how much new sequence is gained when a sequence collection grows. DandD can describe how much human structural variation is being discovered in each new human genome assembly and when discoveries will level off in the future. DandD uses a measure called δ (“delta”), developed initially for data compression. Computing δ directly requires counting k-mers, but DandD can rapidly estimate it using genomic sketches. We also propose δ as an alternative to k-mer-specific cardinalities when computing the Jaccard coefficient, avoiding the pitfalls of a poor choice of k. We demonstrate the utility of DandD’s functions for estimating δ, characterizing the rate of pangenome growth, and computing allpairs similarities using k-independent Jaccard. DandD is open source software available at: https://github.com/jessicabonnie/dandd.

## 1 Introduction

Pangenomes are growing as more high-quality assemblies are produced. Once a sufficient number of assemblies have been added, a pangenome can reach a point of diminishing returns, where each new genome contributes little novel sequence to the collection [1]. Measuring the amount of new sequence per assembly, though, is neither conceptually nor algorithmically straightforward. By surveying studies that computed the average amount of novel sequence per individual in human genome assemblies, Sherman et al [2] found that estimates varied from 0.2 Mbp to 14 Mbp. This wide range in values was attributable to reasonable investigator choices such as selection of alignment parameters or criteria for identifying contigs with novel sequence. In short, the parameter choices led to a wide range of answers, making the end results difficult to compare.

Alignment-free, i.e. *k*-mer based, approaches offer an alternative. Such methods have been used to determine if a pangenome has reached the point of being “closed,” with open/closed status determined by fitting a Heaps-Law model to an empirical *k*-mer growth function [1]. While avoiding many of the parameter-selection pitfalls of alignment-based methods, *k*-mer based approaches, as the name suggests, still require an initial choice of substring (*k*-mer) length, with subsequent measurements dependant on this selection.

We present a new parameter-free method and tool for measuring the amount of sequence in a pangenome based on ideas from string compression. We use a quantity “delta” (*δ*) that measures compressibility of a repetitive string [3]. Other quantities have been proposed for this purpose, including the number of runs in the Burrows-Wheeler Transform (*r*) [4], number of phrases in the Lempel-Ziv parse (*z*) [5], and the size of the string attractor (*γ*) [6]. All these measures have distinct algorithms and interpretations, but *δ* is known to have advantageous bounds compared to the others. For instance, *δ≤γ* for all strings [3].

Computational difficulty among measures quantifying novel sequence varies widely. *z* and *r* require computing a Lempel-Ziv parse or Burrows-Wheeler Transform, respectively, across the entire input string. Computing *γ* is NP complete. Computing *δ*, however, requires little more than a single pass over the input to count *k*-mers. Available tools like KMC[7] can do this efficiently.

Besides being a useful measure of repetitiveness, *δ* is also remarkably easy to estimate. This is true not only when estimating *δ* over a given sequence collection, but also when estimating over *unions* of sequences, as is needed to assess pangenome saturation. Estimating *δ* for sequences and their unions reduces to the problem of estimating set cardinality. Our main insight is that estimating cardinalities over large sequences and their unions is highly efficient using sketches such as MinHash [8, 9] or HyperLogLog [10, 11].

Additionally, we propose a measure called the *k*-independent Jaccard, or KIJ, as an alternative to the Jaccard coefficient. KIJ avoids the risks of preselection of *k*-mer length by using *δ*. Having a principled way to measure similarity without a pre-determined *k* is critically important since poor choices of *k* can lead to incorrect conclusions downstream, as we show in the context of phylogenetic reconstruction.

We describe the algorithms and data structures implemented in the new DandD software tool, which can compute and estimate *δ* in a variety of scenarios relevant to genomics and pangenomics. For example, we demonstrate that (a) DandD can efficiently and accurately estimate *δ* using genomic sketches, (b) DandD can measure how much new sequence is in each new assembled human genome in a pangenome, including when the input consists of assemblies without common coordinates, and finally (c) DandD can be used to compute KIJ, yielding phylogenetic trees that closely match known truth.

## 2 Methods

### 2.1 The delta compressibility measure

Various measures have been proposed for how to quantify the amount of distinct sequence in a pangenome. Some of these measures identified as byproducts of particular compression strategies. For instance, the measure *z* is derived by computing a Lempel-Ziv parse of the pangenome [5]; *z* is equal to the number of phrases in that parse. The measure *r* is obtained by computing the Burrows-Wheeler Transform (BWT) of the pangenome [4]; *r* is equal to the number of maximal same-letter runs in the BWT-transformed string. Other proposals generalize the notion of compressibility to be independent of any compression strategy, such as the string-attractor *γ* [6].

Delta (*δ*) is another measure of compressibility, defined over a pangenome *S*:

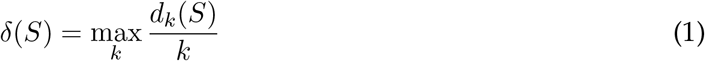

where *d*_*k*_(*S*) is the number of distinct length-*k* substrings among all strings in *S*, i.e. the cardinality of *S*. We use *k*\* to denote the value of *k* that achieves the maximum.

When *S* consists of a single string *s*, the expression *d*_*k*_(*S*)*/k* can be considered to undergo three phases of growth with respect to *k*. For values of *k* that are so short that virtually all possible *k*-mer arrangements of the alphabet appear in *s, d*_*k*_(*S*)*/k* grows exponentially. For values of *k* approaching |*s*|, *d*_*k*_(*S*)*/k* decreases linearly as *k*-mers outgrow *s*, eventually reaching 1 when *k* =|*s*| . For intermediate values of *k*, these trends are in tension; increasing *k* both increases the space of possible *k*-mers, but eventually stops gaining many new distinct *k*-mers. Choosing the *k* that maximizes *d*_*k*_(*S*)*/k* identifies this point of diminishing returns.

*δ* is insensitive to reversals and monotone with respect to appending or prepending symbols to *S*. The *k* in the denominator of Equation 1 links *δ* to other measures. For instance, with *k* in the denominator for *δ*, it is easy to show that *δ≤γ*, where *γ* is the string attractor size [3].

As a function of the substring composition of *S, δ* is comparatively easy to compute relative to other measures. The computation of *z* or *r* is concerned with the entirety of *S*, whereas computing *δ* can be built incrementally by considering *S*’s substrings one-by-one. The advantages of this incremental approach are two-fold: (a) *δ* can be computed simply and in linear time by scanning *S* and counting *k*-mers for an appropriate range of values of *k*, and (b) DNA *k*-mers can be “canonicalized” at the outset, allowing DNA strings and their reverse complements to be treated as equivalent for the purpose of computing *δ*, as is common. During a single scan of *S*, each *k*-mer can be tallied either as itself or as its reverse complement (whichever is lexicographically smaller). For *z* and *r*, allowing for equal treatment of forward and reverse complement strands would require a more drastic approach, e.g. first concatenating *S* with its reverse complement, then running the corresponding algorithm.

### 2.2 Estimating cardinality

*k*-mer counting is resource intensive, potentially requiring a large memory footprint for pangenome inputs. Instead, we propose a method for estimating *δ* by estimating the numerator *d*_*k*_(*S*) (i.e. the cardinality) from Equation 1. The Dashing [11] tool, while chiefly used to estimate similarities between sequencing datasets, can also be used to estimate cardinalities like this via its `dashing card` function, which uses Ertl’s maximum likelihood estimator [11, 12].

Genomic sketches are composable, meaning that sketches built over datasets *A* and *B* can be easily combined to form the sketch for *A∪B*. In the case of HyperLogLog sketches, this is accomplished by simply taking the elementwise maximum of the register values for the sketches of *A* and *B*. The resulting sketch is identical to the one that would have resulted from sketching *A∪B*. Since registers usually number in the thousands-to-millions range, this is a fast operation, using only sequential memory accesses.

### 2.3 Estimating delta

To estimate *δ*, we seek the substring length *k* giving maximal *d*_*k*_(*S*)*/k*, which we denote *k*\*. Determining *k*\* requires a scan, similar to a root-finding procedure. DandD accomplishes this with a simple sweep starting from a user specified initial value of *k*. The sweep tries successively larger and smaller values of *k*, searching for three consecutive values of *k* such that:

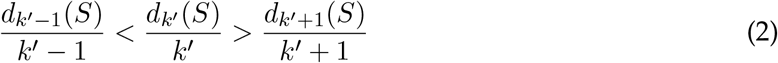

By default, DandD begins its search at *k* = 14. The final value of *k*\* is dataset-dependent, as illustrated in Figure 1, which shows a 0-1 normalized version of *δ* for human, E. coli and salmonella pangenomes.

**Figure 1:**
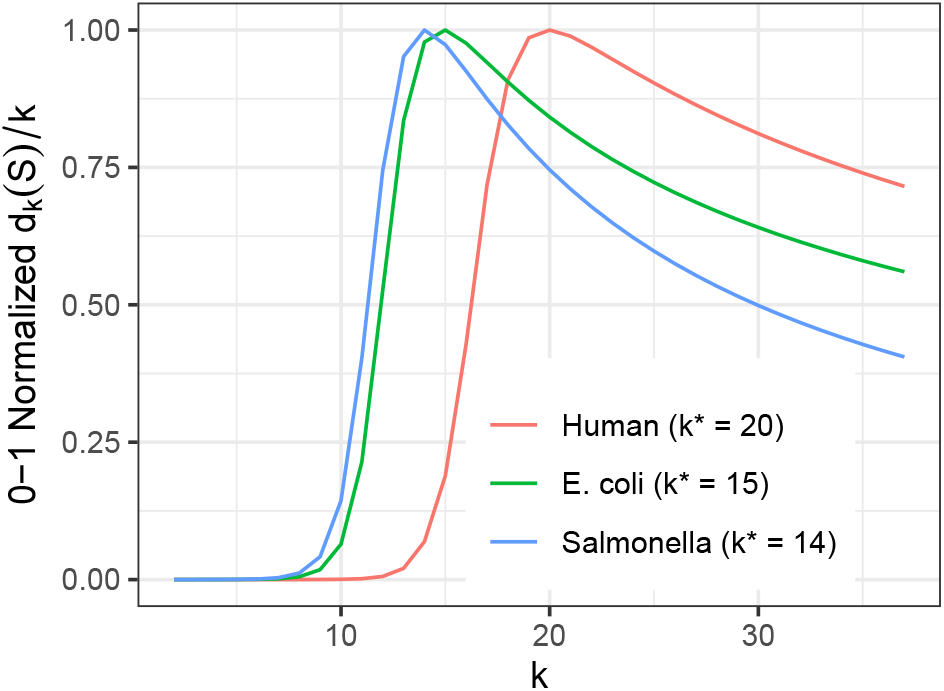
Different pangenomes find their maxima at different values of *k*. Vertical axis shows 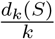 standardized to between 0 and 1 per pangenome. Each pangenome is comprised of multiple distinct genome sequences: H.Sapiens (*n* = 12), E.coli (*n* = 10), and S.enterica (*n* = 10).

In some of DandD’s modes (such as `progressive` and `kij`), it computes *δ* with respect to the union of two or more inputs for which it has previously computed *δ*. In such cases, DandD initializes the search for the union *k*\* by taking the maximum of the previously-computed *k*\**s* of the inputs.

### 2.4 Characterizing sublinear growth

DandD, via the `dandd progressive` command, can characterize the rate of growth of a pangenome by measuring *δ* with respect to cumulative subsets of its constituent genomes. For small pangenomes (up to 6–7 genomes), it can be practical to do examine all possible orderings (permutations) of the genomes. However, for larger real-world pangenomes, it is sufficient to use a random subset of all possible orderings. This method provides a way to estimate the average *δ* for subsets of a given size. That is, by taking the mean of all the values for *δ* obtained after adding the *i*^*th*^ genome in each ordering, we have an estimate for *δ*(all size-*i* subsets).

Past methods for characterizing pangenome growth also made use of random orderings of large collections. These methods seek to determine whether the pangenome is “open” (still accumulating new sequence), or “closed” (substantially complete) [1, 13, 14]. DandD provides a new way of performing this analysis over genome sequences in a parameter-free way, not requiring foreknowledge of where genes are located or how to choose an appropriate value of *k*.

The `dandd progressive` command allows the user to provide a set of genomes and a desired number of orderings to try. DandD will then (1) preprocess all of the inputs individually, (2) generate the random orderings, and then (3) iterate through each ordering, “progressively” building larger unions by accumulating one more genome at each step. The `dandd progressive` command outputs a file describing, for each step of each ordering, which genome was added in that step and the value of *δ* for the new union.

Besides giving useful plots (seen in Results 3.3 below), this output can be used to fit a Heap’s Law model to the change in delta (Δ*δ*) at each step. After fitting, the fit value of the Heaps-Law *α* parameter can be used to characterize whether the pangenome is open (*α ≤* 1) or closed (*α >* 1).

### 2.5 K independent Jaccard

The Jaccard coefficient is a widely used metric for comparing large datasets:

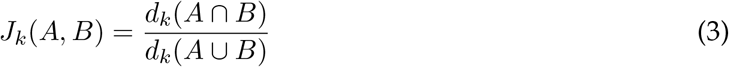

For consistency we use *d*_*k*_(*A*) (rather than |*A*| _*k*_) to denote the cardinality of the set of *k*-mers in text *A*. Methods like MinHash estimate this quantity directly. Methods based on the HyperLogLog sketch, like Dashing, obtain separate cardinalities *d*_*k*_(*A*), *d*_*k*_(*B*) and *d*_*k*_(*A∪B*) and compute *J*_*k*_ using an expression equivalent (by the inclusion-exclusion principle) to the one above:

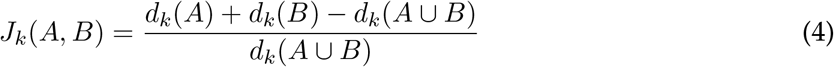

The above expressions have *k*-mer length *k* as a parameter. To obtain a *k*-independent notion of Jaccard coefficient, we replace *d*_*k*_ with *δ*:

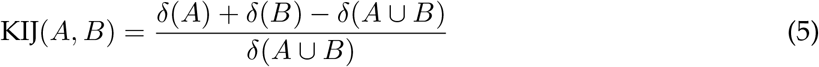

Following this formula, the task of computing or estimating KIJ reduces to the task of obtaining *δ*(*A*), *δ*(*B*) and *δ*(*A∪B*) (or estimates thereof). DandD provides a command (`dandd kij`) to compute all-pairwise KIJs given two or more input FASTA files. Since downstream tools may expect to receive distances rather than similarities, DandD can output all-pairwise 1-KIJs instead.

### 2.6 Caching and lazy evaluation of sketches

The most time and memory intensive step of solving for *δ* is the creation of the component sketches from the input FASTA files. Computationally, it is far simpler to produce the union of two or more sketches or estimate cardinality from existing sketches. Given these uneven resource requirements, DandD is designed to reuse sketches within tasks and across experiments within the same pangenome.DandD reduces its footprint and prevents the production of duplicate sketches by caching sketches on the filesystem and tracking the sketches it has already built. To maintain the association between a union sketch and its component FASTAs, the sketch file is named using a checksum over the constituent FASTAs as computed by the cryptographic hash function BLAKE. This serves a dual purpose of insuring that each combination of input FASTAs is sketched only once, regardless of order, and providing a mechanism to confirm agreement between the currently available FASTAs and any information pertaining to them that may be stored within DandD’s tree structure. In addition to a naming convention, DandD also creates a directory structure to store intermediate sketches and databases for easy reuse and access.

When the user specifies the same sketch directory across many invocations of DandD, they will get the maximum benefit of reuse of sketch files overall.

### 2.7 Exact mode

DandD includes an “exact” option (`--exact` option) which enables computation of *δ* directly by way of *k*-mer counting. This option can be used in combination with any of DandD’s modes (`tree`, `progressive`, `kij`, etc). Instead of using Dashing, “exact” mode uses KMC3 for counting (via the `kmc` command) and unioning (via `kmc tools complex`) [7]. Just as Dashing must build a sketch prior to estimating cardinalities, KMC3 builds a “database” of *k*-mer counts.

Note that while unioning two Dashing sketches requires only an elementwise maximum over the sketches, unioning two KMC databases requires a merge sort over all the *k*-mers and counts. This can be quite expensive, as detailed in Table 1.

**Table 1:**
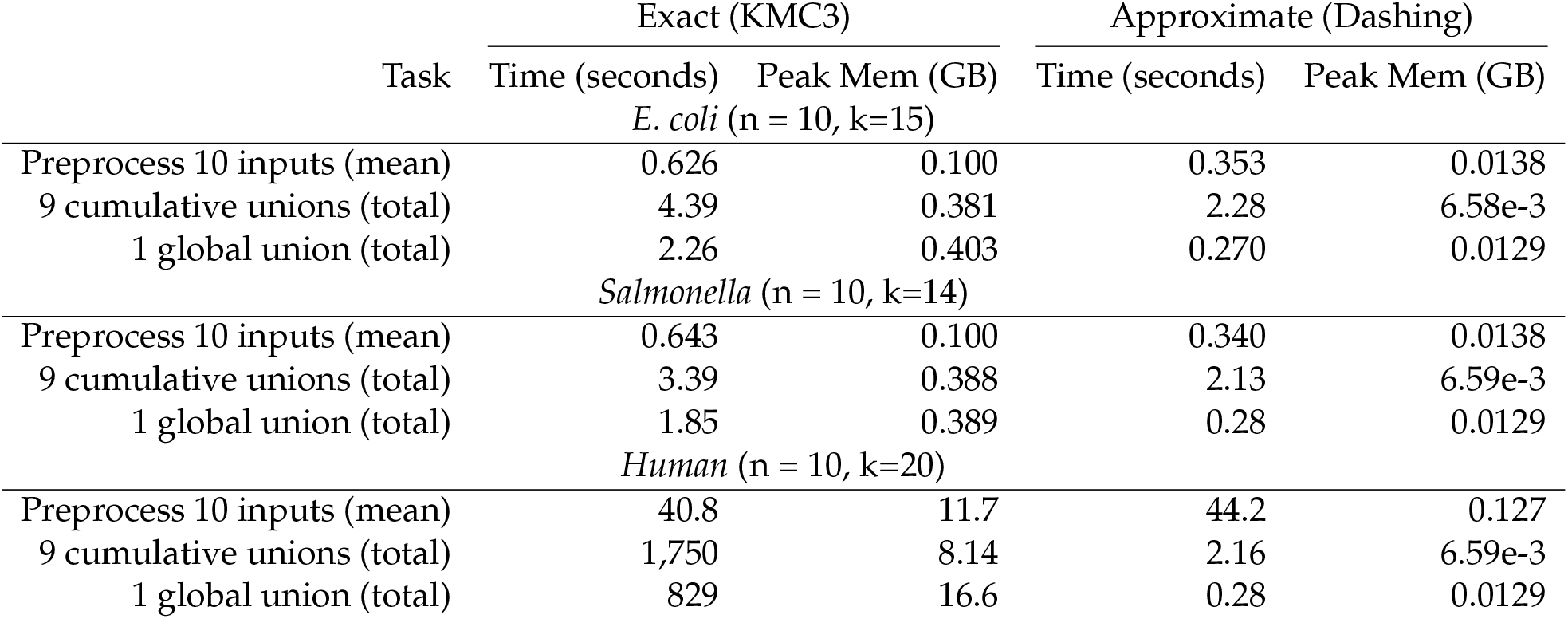
Computational efficiency of KMC3 versus Dashing for preprocessing and unioning. For KMC3, preprocessing consists of building *k*-mer count databases. For Dashing, it consists of building genomic sketches. We analyzed three collections of genome assemblies: E. coli (n=10, k=15), salmonella (n=10, k=14), and human (n=10, k=20). For simplicity, we chose a single *k* for each which was appropriate to the species. We measured the time and memory (resident set size) required to preprocess the 10 inputs on average, reported in rows labeled “Preprocess 10 inputs.” We chose a random ordering of the genomes and measured the resources required to perform a series of unions, each adding one additional genome to the union (“9 cumulative unions”). We also measured resources required to union all preprocessed datasets at once (“1 global union”).

DandD’s file naming scheme together with the metadata it saves allows users to locate KMC databases and Dashing sketches corresponding to particular inputs and their unions. This facilitates further experimentation; i.e. a user can use KMC3’s tools to explore which *k*-mers contributed to a particularly large increase in *δ*.

## 3 Results

### 3.1 Behavior of *δ*, r and z in practice

We began with an empirical study of the relationship between *z, r* and *δ* using real genome sequence data. We collected 50 *Salmonella enterica* genomes from Refseq, putting them in an arbitrary order. We computed *z, r* and *δ* for sets of these genomes, starting with the first and cumulatively adding one at a time. In the case of *z*, we used a combination of the `newscanNT.x` tool from Big-BWT [15] and the `lz 77` tool from PFP LZ77 [16]. To compute *r*, we used the `pfbwt-f64` tool [15]. For *δ*, we used DandD in its `--exact` mode, which in turn uses KMC3 [7]. For *r* and *z*, both the genomes and their reverse complements were added to the set at the same time. For *δ*, we accomplished this through *k*-mer canonicalization.

Results are shown in Figure 2, with the three measures normalized to a range of 0 to 1. There is an obvious, strong relationship between the measures, with no two of the normalized measures differing by more than *±* 0.0273 at any point.

**Figure 2:**
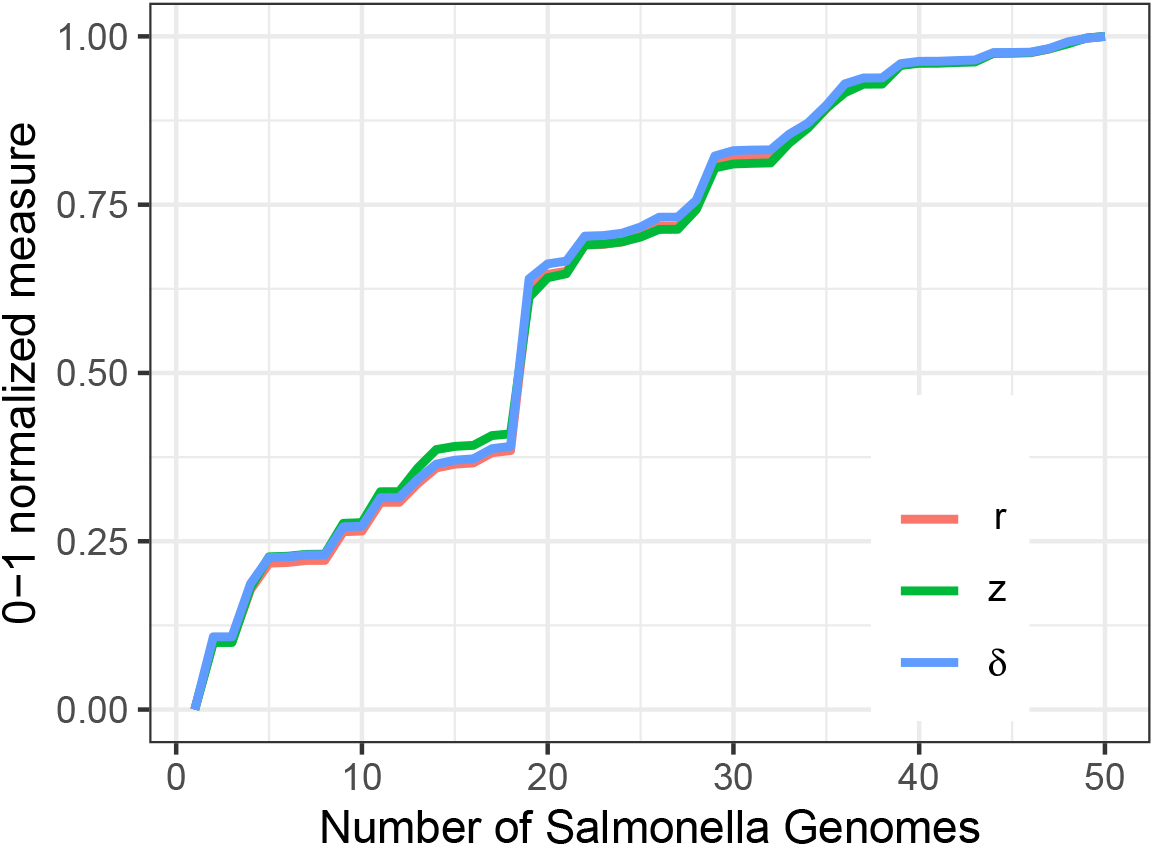
*δ, r*, and *z* varying together.

Figure A1 (Appendix) also shows the wall clock time required by these methods for computing *z, r* and *δ*, with the KMC3-based method for computing *δ* being the fastest.

### 3.2 Efficient cardinality estimation with Dashing

We sought to compare the computational performance and accuracy of DandD’s two modes, the`--exact` mode (“exact”) and the sketch-based mode (“approximate”). We ran DandD in both modes on three datasets: 10 E. coli genomes, 10 salmonella genomes, and 12 human genomes.

We performed all the computations for a single value of *k*. These methods are being assessed for performance in computing (or estimating) the cardinatlity *d*_*k*_(*S*), not *δ*. Given that computing *δ* means repeatedly performing this process for several values of *k*, this provides sufficient basis for comparing the methods.

For each dataset and method, we performed three computations. First, we used the selected method to build a *k*-mer database (for “exact”) or sketch (for “approximate”) over each input genome. We used the selected method to perform a series of cumulative unions, starting from a single database/sketch and combining them in successive steps until all genomes are included. Finally, we performed a single global union of all of the individual databases/sketches. In all cases, we used `/usr/bin/time -v` to measure the time and the maximum memory usage.

In nearly all cases, the “approximate” method is faster than the “exact” method (Table 1). This is particularly true for the union steps for the human genome inputs, where the approximate method is as much as three orders of magnitude faster than the exact method. The “approximate” method’s peak memory footprint is consistently smaller than the exact method’s, sometimes by two orders of magnitude.

Further, for every experiment described in Table 1, we used Dashing to estimate the cardinality from the sketch produced and compared this to the true cardinality as computed by KMC3. We computed the relative error of the Dashing estimate as |*D − K*|*/K*, where *D* is Dashing’s estimate and *K* is KMC3’s exact count. Overall, the mean relative error was 6.537 *×* 10^*−*4^. The maximum observed relative error was 1.32 *×* 10^*−*3^.

### 3.3 Using DandD to characterize sublinearity and openness

We ran the `dandd progressive` command on a set of 34 human haplotypes taken from the Human Genome Structural Variation Consortium (HGSVC2) project [17]. The haplotypes were chosen to all have an X chromosome, to avoid large increments in *δ* due only to the addition of the Y chromosome.

HGSVC2 is organized into VCF files for each distinct variant type: single nucleotide variants (SNVs), small insertions and deletions (Indels), or structural variants (SVs). We repeated our experiment across different subsets of variant catgories. For instance, to create the FASTA sequences used as input to our “SNV only” experiment, we used `bcftools consensus` to create 34 haplotype-specific FASTA files, taking rows from the VCF file containing SNVs. For the “SNVs + indels” experiment, we did the same but taking rows from both the SNV and Indel VCF files. For the “SNVs + SVs + indels” experiment, we did the same but taking rows from all three VCFs.

In all cases, the individual haplotype FASTA were then provided to `dandd progressive`. Since it is impractical to attempt all orderings of 34 haplotypes, we used DandD’s `--norder 120` option to randomly generate a series of 120 possible permutations.

We performed a corresponding experiment for 56 haplotypes from the Human Pangenome Reference Consortium (HPRC) [18], taken from 28 individuals annotated as female. In this case, the FASTA inputs to DandD were the phased assemblies provided by the HPRC. There is no accompanying VCF file describing variants or variant types, giving us no way to stratify by variant type as we did for HGSVC2. But DandD is applicable regardless, since it simply accepts any FASTA inputs. We used DandD’s `--norder 90` option to randomly generate a series of 90 possible permutations, 5 of which are shown in Figure 3.3B.

**Figure 3:**
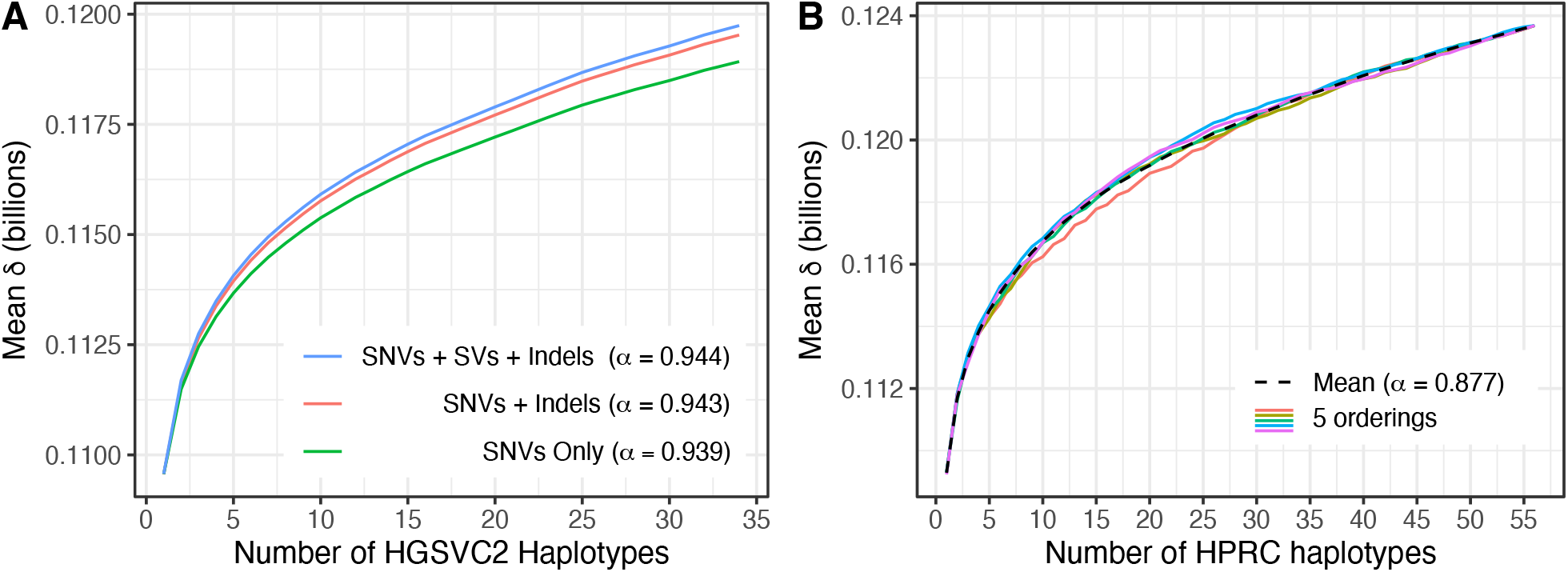
Empirical growth of *δ* for human haplotypes from the HGSVC2 project (panel A) and the HPRC (B). For (A), lines show mean value of *δ* for each genome count across all 120 orderings. Colors denote how the variants were subsetted before constructing the FASTA files given to DandD. E.g. the blue line corresponds to genomes edited to include SNVs, small indels and structural variants, whereas the green line corresponds to genomes edited to include only SNVs. For (B), the dotted line shows the mean value of *δ* at each genome count across all 90 orderings. The colored lines show the particular values of *δ* from a random subset of 5 orderings.

In all cases, we computed a Heaps-Law fit to the mean *δ* values and recorded *α* for each. Fit values of *α* are reported in the Figure 3.3 legends.

As seen in Figure 3.3, all experiments showed sublinear growth in mean *δ*. For HGSVC2, we observed that “SNVs + SVs + Indels” had the highest *δ* overall, with “SNVs + Indels” having slightly lower values, and “SNVs only” having still lower values. This is expected, since the inclusion of each additional variant class should lead to new distinct sequence. The Heaps-Law *α* was approximately the same for all three HGSVC2 variant subsets, *≈* 0.94 in all cases. Thus, all variant classes lead to the same conclusion about “openness” of the HGSVC2 pangenome, i.e. *α*≤ 1 indicates it is open.

The HPRC data also showed sublinear growth, and a range of *δ*’s quite similar to those observed for HGSVC2. The Heaps-Law *α* = 0.873, again indicating an open pangenome. The fact that *α* is lower for HPRC may indicate that the *de novo* assemblies from long reads give access to a wider array of genetic variants, which in turn requires more genomes to saturate the pangenome. However, this is hard to disentangle from the effects of sequencing errors, which can be counted (spuriously) as novel sequence. Removing the effect of sequencing errors is an important problem, which we return to in the Discussion.

### 3.4 Evaluating KIJ

To evaluate the utility of the *k*-independent Jaccard (KIJ) measure, we used benchmarking datasets from the AFproject [19]. These benchmarks use real data to evaluate the quality of clusterings created by alignment-free methods. In particular, we used the 25 fish mitochondrial genomes and the 29 E. coli genomes.

For KIJ, we computed all pairwise distances between the sequences in the dataset. Since KIJ measures similarity, we report 1-KIJ as the distance. From this point, we proceeded with the steps of the AFproject protocol, which constructs a tree from the pairwise-distances, then compares that true to a curated tree. The ultimate result is a normalized version of the Robinson-Foulds distance (nRF), which measures the degree of structural difference between two trees having the same set of sequences at the leaves. A low nRF indicates that the distances provided reflected the true phylogenetic relationships between the sequences.

Having done this for 1-KIJ distances, we repeated the process for distances based on *k*-specific Jaccard coefficients for a range of *k*s. We computed *J*_*k*_ and reported a matrix of pairwise 1*− J*_*k*_ distances for *k* = 2 to 59. nRFs obtained for each of these are shown in Figure 4.

**Figure 4:**
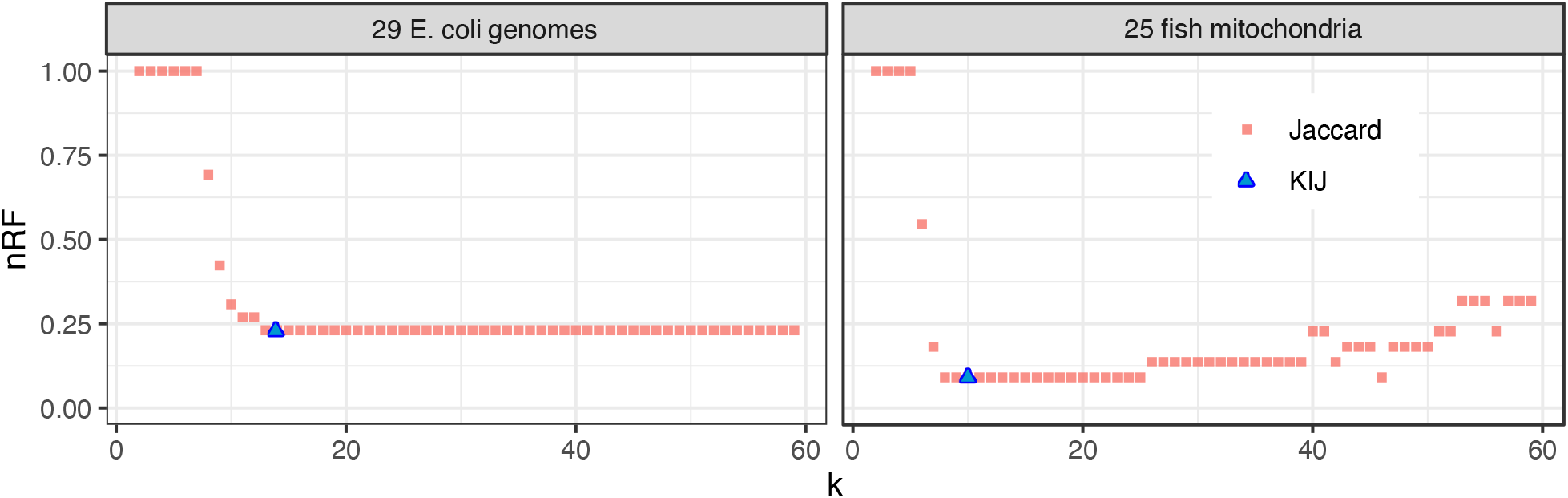
Clustering based on KIJ distances achieves low normalized Robinson-Foulds (nRF) distance with respect to the true phylogeny compared to clusterings based on the typical Jaccard coefficient.

We observed that the ability of the 1*− J*_*k*_ distance to produce a accurate predicted tree is dependent on the choice of *k*. A too-small value of *k* leads to non-specific distances that cannot distinguish the phylogenetic relationships, leading to high nRF toward the left-hand side of the plots in Figure 4. A too-large value of *k* can deplete the number of common *k*-mers between related sequences in a way that obscures their relationship, as seen toward the right-hand side of the “25 fish mitochondria” plot in Figure 4, where nRF climbs after *k* grows past 25.

The 1-KIJ distance, on the other hand, strikes a balance between these extremes. In both cases, the 1-KIJ distance achieves minimal nRF compared to all of the 1-*J*_*k*_ distances.

## 4 Discussion

As sequencing technology improves, new genome assemblies will arrive more quickly. It will be increasingly important to identify when a collection of genomes has reached a point of saturation, i.e. when it represents a taxonomic grouping in a complete fashion without excess accumulation of rare variation. Since pangenomes can be seen as collections of repetitive strings, theory concerning compressibility provides tools well suited to this problem.

Building on the *δ* measure, DandD provides an efficient and interpretable way to measure the growth of pangenomes and compare large sequence collections. *δ* has theoretical advantages but is also remarkably easy to compute. Genomic sketches make *δ* particularly easy to estimate over pangenomes and their unions. Further, *δ* provides a parameter-free way of quantifying the amount of distinct sequence in a pangenome, sidestepping any dependence on parameters.

The methods underlying DandD treat pangenomes as sets, with KIJ providing a quantification similar to the Jaccard coefficient between sets. However, another way to represent and sketch pangenomes would be as multisets, where each item (i.e. *k*-mer) has an associated count; e.g. the number of times it occurs in the pangenome. Genomic sketches like SetSketch can accommodate counts, and Dashing 2 [20] can compute probability-weighted version of the Jaccard coefficient. In the future, it will be important to evaluate whether consideration of counts can be naturally combined with these compressibility measures.

It should be possible to convert the 1 *−* KIJ distance measure discussed in Results 3.4 into a Mash distance [9], though with the additional complication that KIJ is a function of three separate *δ* measures, *δ*(*A*), *δ*(*B*) and *δ*(*A∪B*). These may use different underlying choices for *k*\*, creating ambiguity in how *k* should be specified in the Mash distance formula. In the future, it will be important to study how to handle multiple distinct *k*’s in the Mash distance formula, and to evaluate how Mash distances derived from KIJ perform relative to those derived from *J*_*k*_.

## 5 Author contributions

JKB, OA and BL conceived the method, designed the experiments and ran the experiments. JKB and BL wrote the paper. JKB wrote the DandD software tool. JKB, OA and BL edited and approved the final manuscript.

## 6 Acknowledgments

We thank Aaron Quinlan and Dominik Kempa for helpful discussions.

## 8 Availability of data and materials

Open source source code for the DandD software is at

https://github.com/jessicabonnie/dandd

Scripts for performing the experiments described in this manuscript are at

https://github.com/jessicabonnie/dandd_experiments

Lists of the accessions for each of the tables and figures in this manuscript are provided at

shttps://github.com/jessicabonnie/dandd_experiments/tree/main/accessions

## Appendix

**Figure A1:**
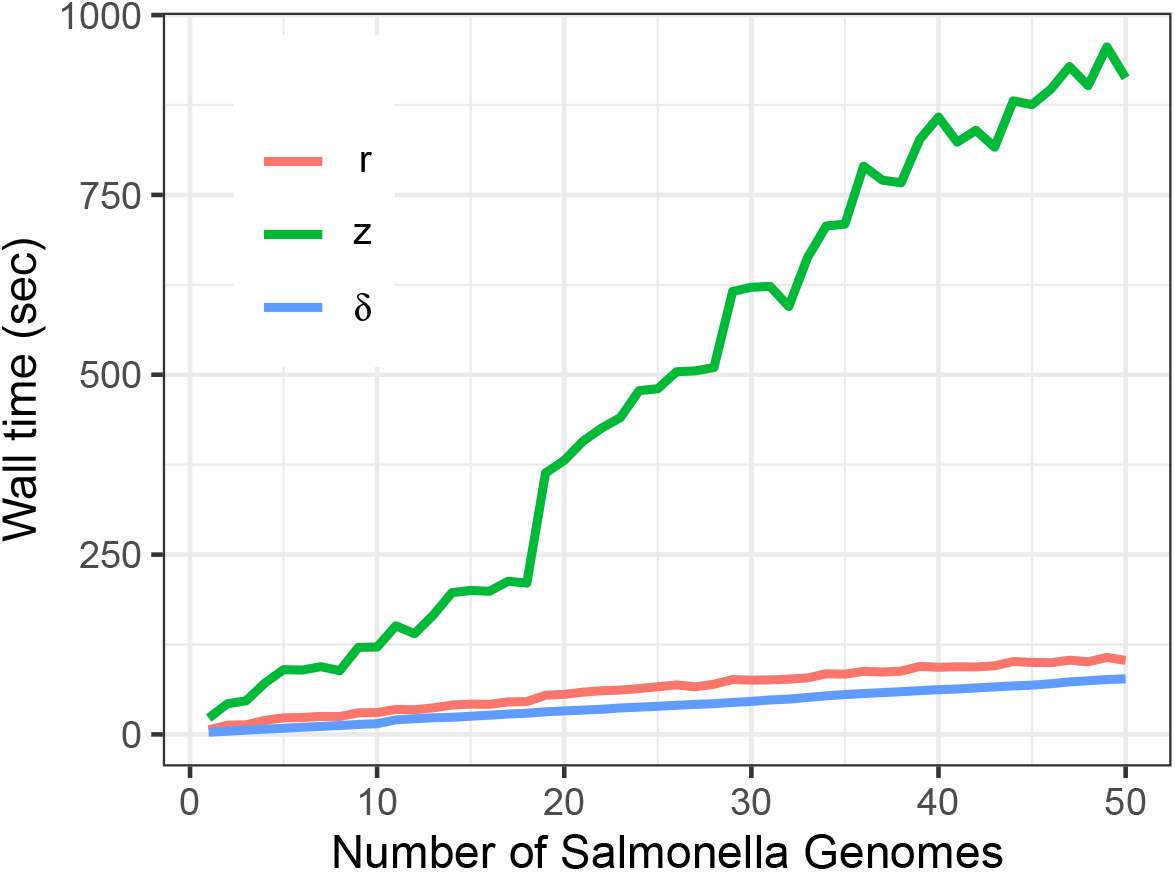
Wall clock time required to compute *δ, r*, and *z*. All experiments used a single thread of execution. We note that these benchmarks use a particular set of tools to compute each measure. It does not necessarily convey the inherent difficulty of computing the measures. A related caveat is that *z* is computed using a series of tools that first computes a prefix-free parse, from which *r* can be immediately inferred. In this way, the method we used to estimate *z* was bound to use more wall time than the method used to estimate *r*. A method that computes *z* directly, not by way of the prefix-free parse, might be more performant.

